# Antiviral activities of two nucleos(t)ide analogs against vaccinia and mpox viruses in primary human fibroblasts

**DOI:** 10.1101/2023.03.23.533943

**Authors:** Lara Dsouza, Anil Pant, Samuel Offei, Lalita Priyamvada, Blake Pope, Panayampalli S. Satheshkumar, Zhengqiang Wang, Zhilong Yang

**Author notes:** Correspondence (SPS); (ZW); (ZY).

## Abstract

Many poxviruses are significant human and animal pathogens, including viruses that cause smallpox and mpox. Identification of inhibitors of poxvirus replication is critical for drug development to manage poxvirus threats. Here we tested two compounds, nucleoside trifluridine and nucleotide adefovir dipivoxil, for antiviral activities against vaccinia virus (VACV) and mpox virus (MPXV) in physiologically relevant primary human fibroblasts. Both trifluridine and adefovir dipivoxil potently inhibited replication of VACV and MPXV (MA001 2022 isolate) in a plaque assay. Upon further characterization, they both conferred high potency in inhibiting VACV replication with half maximal effective concentrations (EC_50_) at low nanomolar levels in our recently developed assay based on a recombinant VACV secreted Gaussia luciferase. Our results further validated that the recombinant VACV with Gaussia luciferase secretion is a highly reliable, rapid, non-disruptive, and simple reporter tool for identification and chracterization of poxvirus inhibitors. Both compounds inhibited VACV DNA replication and downstream viral gene expression. Given that both compounds are FDA-approved drugs, and trifluridine is used to treat ocular vaccinia in medical practice due to its antiviral activity, our results suggest that it holds great promise to further test trifluridine and adefovir dipivoxil for countering poxvirus infection, including mpox.

## Introduction

The family *Poxviridae* comprises 22 genera with 83 species based on the 2021 International Committee on Taxonomy of Viruses (ICTV) release. They cause a broad range of human and animal diseases. The orthopoxvirus genus contain 12 known species including high consequence human pathogens, such as variola virus that causes smallpox, and mpox/monkeypox virus (MPXV). Historically, smallpox had been accounted for the most human lives among all infectious diseases. It is estimated that ∼300 million people died of smallpox in the first 80 years of the 20^th^ century alone before its eradication in 1980 ^1^. Despite its eradication^2^, potential smallpox re-emerging from unsecured stocks or by synthetic biology approach remains a major national security concern ^3,4^, particularly due to the rapid decline of population immunity against smallpox after the cease of smallpox vaccination. The loss of the cross-protection by immunity against smallpox also increases the danger of other orthopoxvirus infections. Consequently, other zoonotic orthopoxviruses may emerge to pose significant threats to public health ^5^. This is exemplified by the ongoing global mpox outbreak with over 85,000 reported cases in more than 110 countries (∼30,000 in the USA). The global outbreak of mpox also underlines the pandemic potential of MPXV, of which future outbreaks are expected^6,7^.

Other orthopoxviruses may also emerge to infect humans. For example, a novel orthopoxvirus caused infections in four human individuals in Alaska in recent years, which is believed to have been transmitted by animals ^8,9^. Such orthopoxviruses may evolve to adapt human host over time and cause more serious concerns.

FDA has approved two drugs for strategic stockpiling against smallpox: tecovirimat^10^ (**1, Fig. 1**) and brincidofovir^11^ (BCV, **2, Fig. 1**), the lipid prodrug of nucleotide analog cidofovir (**3, Fig. 1**). However, the clinical efficacy of BCV against mpox is not promising ^12,13^, and the clinical use of cidofovir for treating human cytomegalovirus is associated with severe adverse effect^14-16^ and drug resistance^17-19^. With a distinct mechanism of action, tecovirimat inhibits viral release by targeting the viral extracellular envelop protein VP37^20,21,22^. Although tecovirimat has shown promising efficacy in some mpox cases^23^, the clinical data are still very limited ^24,25^. Importantly, tecovirimat has a low barrier to viral resistance based on studies from literature^22^ and FDA released data^26^, and resistant mutants are expected after extensive use. Therefore, it is critically important to develop new chemical entities against orthopoxviruses to provide valuable leads for rapid and effective countermeasures against re-emerging smallpox, mpox outbreaks, and other emerging orthopoxviruses.

**Fig. 1.**
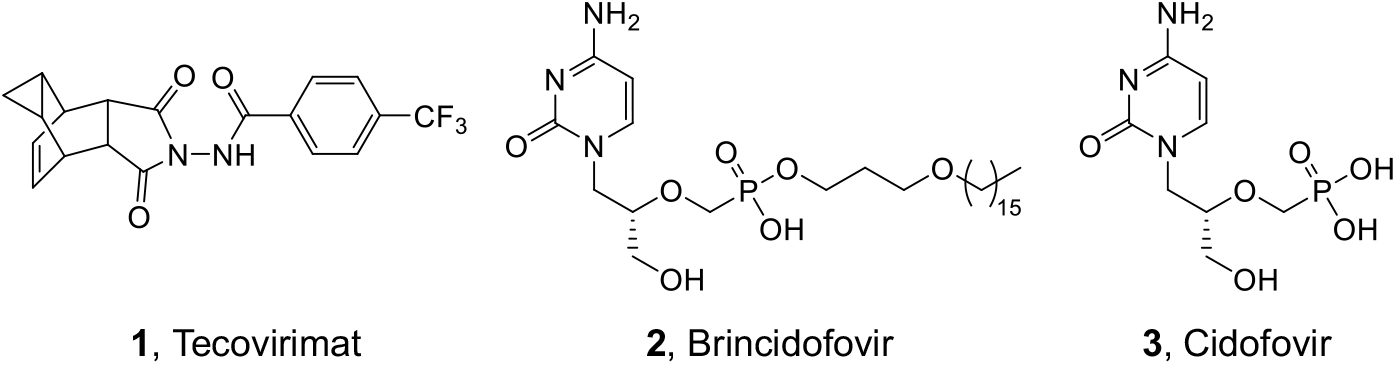
Structures of FDA-approved smallpox drugs. Tecovirimat (**1**) inhibits viral release by targeting the envelop protein VP37. Brincidofovir (**2**) is the prodrug of Cidofovir (**3**) which targets viral polymerase.

Vaccinia virus (VACV) is the prototype poxvirus and is a highly relevant surrogate to study high pathogenic poxviruses (e.g., mpox and smallpox viruses) due to their high similarity with >95% genome identity ^27^. Using VACV, we previously screened a library comprising FDA-approved antiviral drugs and a Selleck bioactives, and identified many hits of VACV inhibitors ^28^. Here we further characterized antiviral activities of two nucleos(t)ide analogs: trifluridine and adefovir dipivoxil. Significantly, they both potently inhibited VACV and MPXV replication in physiologically relevant primary human foreskin fibroblasts (HFFs) without cytotoxicity. Mechanistically, they both target the DNA replication stage of viral infection. Our findings provide two strong anti-poxvirus hits for further development.

## Results

### Trifluridine and adefovir dipivoxil potently inhibit VACV replication in primary HFFs

We have previously screened focused compound libraries for VACV inhibitors using a reporter VACV expressing Gaussia luciferase (Gluc) (vLGluc) in transformed HeLa cells, and have identified a number of strong hits ^28^, including several nucleos(t)ide analogs. To further confirm the antiviral effects of trifluridine (**4, Fig. 2A**) and adefovir dipivoxil (**5, Fig. 2A**), we tested their effects on VACV replication in primary HFFs using plaque assay, the gold standard method of infectious viral yield measurement. It is worth noting that dermal fibroblasts are physiologically relevant to orthopoxvirus infection as they are among the major cells in poxvirus infection and dissemination ^29^. Under a multiplicity of infection (MOI) of 0.01 and 48 h infection incubation time, trifluridine and adefovir dipivoxil strongly suppressed VACV yield by ∼9,500- and 4,500-fold, respectively, at 10 µM (**Fig. 2B**), without reducing viability of HFFs (**Fig. 2C**). As a positive control, cytarabine (AraC, **6, Fig. 2A**), a well-studied compound that block VACV genome replication, also strongly suppressed VACV replication in HFFs by 24,000-fold (**Fig. 2BC**).

**Fig. 2.**
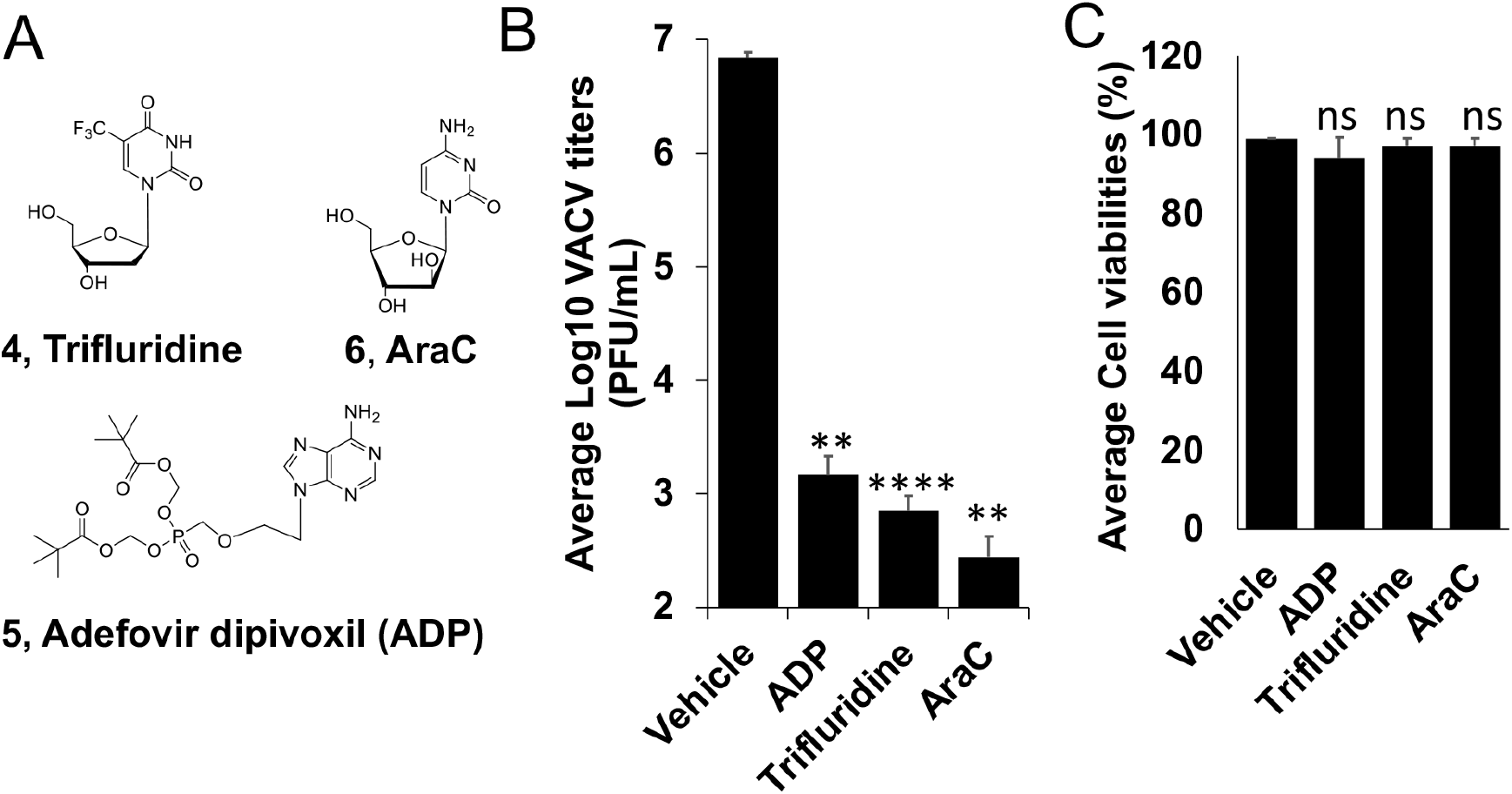
Inhibition of VACV replication by trifluridine and Adefovir dipivoxil (ADP) by plaque assay. **(A)** Structures of trifluridine (4), adefovir dipivoxil (ADP, 5), and AraC (6). (**B**) HFFs were infected with VACV at an MOI of 0.01 treated with indicated compounds at 10 µM for 48 h. Viral yields were titrated using a plaque assay on BS-C-1 cells. (**C**) HFF cell viabilities were measured 48 h after the cells were treated with indicated compounds at 10 µM. At least three repeats were performed for all assays. **0.001<p ≤ 0.01, ns, not significant.

We next determined the half maximal effective concentration (EC_50_) of trifluridine and adefovir dipivoxil in inhibiting VACV replication in HFFs using the vLGluc. The vLGluc is a recombinant VACV expressing Gluc under a viral later promoter used in our initial screening of VACV inhibitors ^28,30^. We first tested to see if this assay is suitable to measure VACV inhibitor’s EC_50_ using BCV ^31^. With an MOI of 0.01 and 24 h infection incubation time, the EC_50_ of BCV was determined to be ∼38 nM in HFFs (**Fig. 3AC**), which is similar to the reported values ^32,33^, suggesting that vLGluc is suitable for EC_50_ measurement. Using the same MOI and incubation time, we determined the EC_50_s of trifluridine (EC_50_ =138 nM) and adefovir dipivoxil (EC_50_ = 302 nM) (**Fig. 3AC**). The EC_50_ of AraC was also determined (EC_50_ =123 nM) (**Fig. 3A**). Remarkably, both trifluridine and adefovir dipivoxil, as well as the positive control compounds BCV and AraC, caused no significant cytotoxicity in HFFs at high concentrations (CC_50_ > 250 µM for trifluridine, AraC, and adefovir dipivoxil, CC50>50 µM for brincidofovir) as measured in an MTT assay assessing the metabolic activity of the cells (**Fig. 3BC**).

**Fig. 3.**
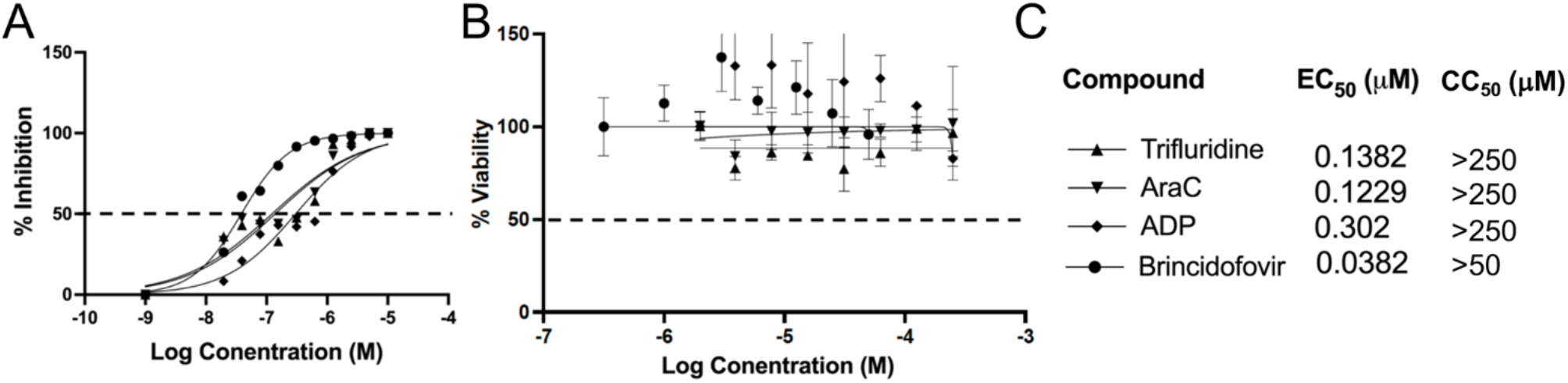
Measurement of EC_50_ and CC_50_ of indicated compounds in HFFs. **(A)** HFFs were infected with vLGluc at an MOI of 0.01 and treated with indicated individual compounds at a series of concentrations (or vehicle DMSO) for 24h. Gluc activities were measured to determine the EC_50_. (**B)** HFFs cell viabilities were determined by an MTT assay after incubated with indicated compounds at a series of concentrations for 24 h. **(C)** EC_50_ and CC_50_ of the compounds in A and B are shown. At least three repeats were performed for all experiments.

Together, the above results establish trifluridine and adefovir dipivoxil inhibit VACV with EC_50_s at low nM low cytotoxic effects in HFFs. Our results also further validate the Gluc expressing reporter VACV as a valuable tool in poxvirus inhibitor identification and chracterization.

### Trifluridine and adefovir dipivoxil inhibit VACV genome replication

Poxvirus replication are divided into the following steps: entry, early gene expression, uncoating, DNA replication, intermediate gene expression, late gene expression, and post gene expression events such as viral morphogenesis, assembly, and spreading ^34^. As nucleos(t)ide analogs, trifluridine and adefovir dipivoxil presumably inhibit VACV replication mainly at the DNA replication stage, although the post DNA replication gene expression may also be affected due to the excessive need of RNA synthesis. To test this hypothesis, we first used recombinant VACVs with stage-specific Gluc reporter genes. In addition to the VACV encoding GLuc under the late F17R promoter (vLGluc), two other recombinant VACVs were also used: in one, the GLuc gene is under the control of the VACV early C11R (vEGluc) promoter, and in the other it is under control of the G8R intermediate (vIntGluc) promoter ^30^. The C11R, G8R and F17R genes are well-characterized, exclusively early, intermediate and late VACV genes, and their promoters can be used to effectively distinguish stages of VACV gene expression^35,36^. Neither trifluridine nor adefovir dipivoxil affected the Gluc expression under the VACV early C11R promoter (**Fig. 4A**), while they both strongly inhibited Gluc expression under intermediate G8R and late F17R promoters (**Fig. 4BC**). The trends were highly similar to AraC treatment (**Fig. 4A-C**).

**Fig. 4.**
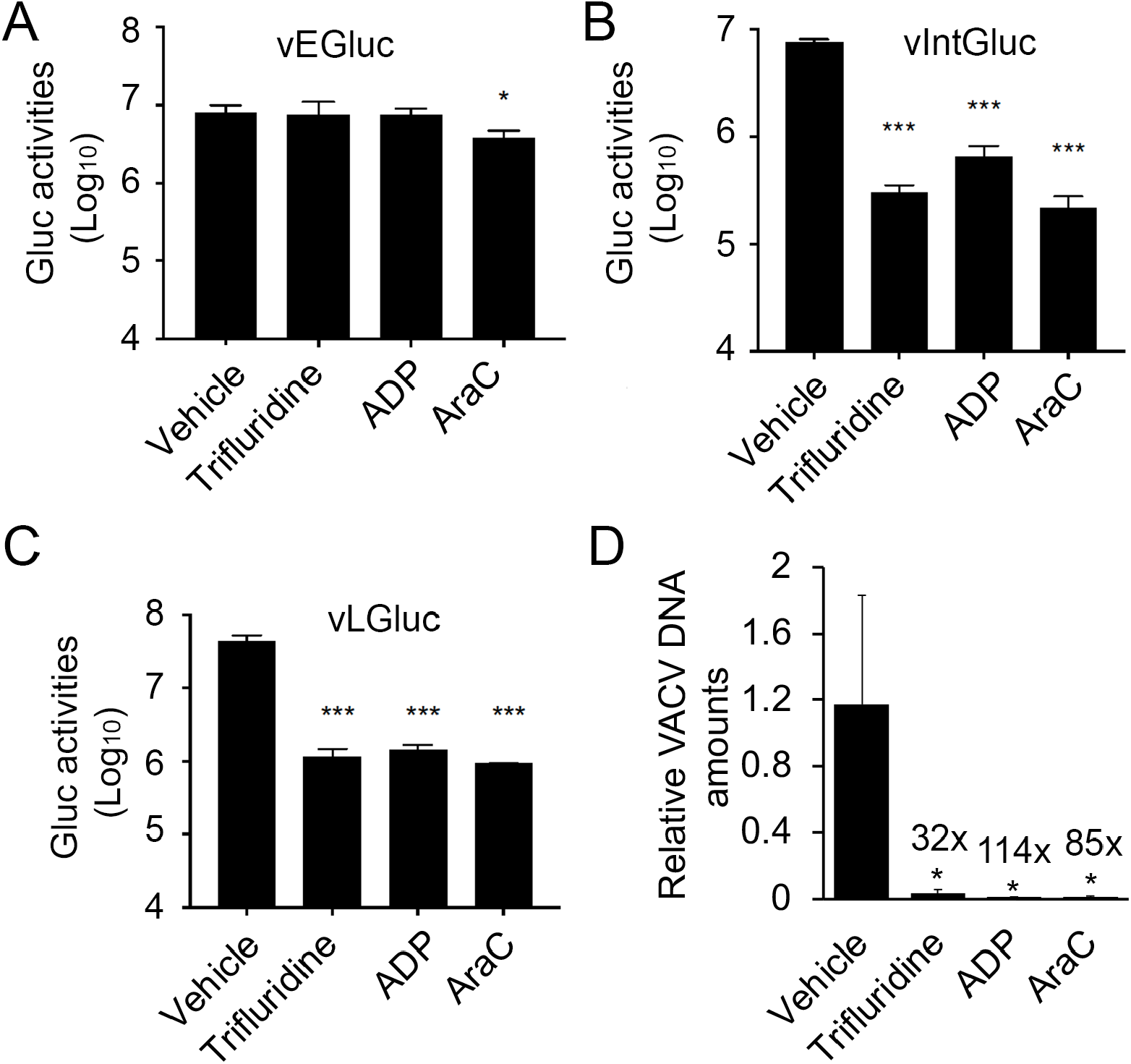
Trifluridine and adefovir dipivoxil suppress VACV DNA replication and post-replicative gene expression, but not early gene expression. **(A-C)** HFFs were infected with vEGluc (**A**), vIntGluc (**B**), and vLGluc (**C**) at an MOI of 2 and treated with indicated compounds at 10 µM, respectively, or vehicle DMSO. Gluc activities were measured at 4 h (vEGluc), 8 h (vIntGluc), and 8 h (vLGluc), respectively. (**D)** HFFs were infected with VACV at an MOI of 2 in the presence of indicated compounds at 10 µM for 8 h. relative amounts of Viral DNA were determined by real-time PCR using VACV specific primers. At least three repeats were performed for all assays. *0.01<p ≤ 0.05; ***0.001<p ≤ 0.01; ns, not significant.

As poxvirus intermediate and late gene expression is dependent on viral genomic DNA replication, we then examined VACV DNA levels in the presence or absence of individual compounds. We found that trifluridine, adefovir dipivoxil, and the positive control AraC, strongly reduced viral DNA levels by 32- to 114-fold (**Fig. 4D**), respectively. Together, these results confirmed that trifluridine and adefovir dipivoxil mainly function to restrict VACV DNA synthesis.

### Trifluridine and adefovir dipivoxil significantly inhibit MPXV replication in primary HFFs

We used two methods to examine the effects of trifluridine and adefovir dipivoxil on MPXV replication. In one method, we used a WA strain MPXV-USA-2003-044 expressing firefly luciferase (Fluc) under a viral early/late promoter (luc+ MPXV) gene as the reporter. We observed that trifluridine and adefovir dipivoxil strongly inhibited MPXV replication with similar potency to AraC (**Fig. 5A**). We also tested the inhibitory effects on an MPXV-MA001 2022 isolate using a plaque assay and found that both trifluridine and adefovir dipivoxil significantly suppressed MPXV replication (**Fig. 5B**).

**Fig. 5.**
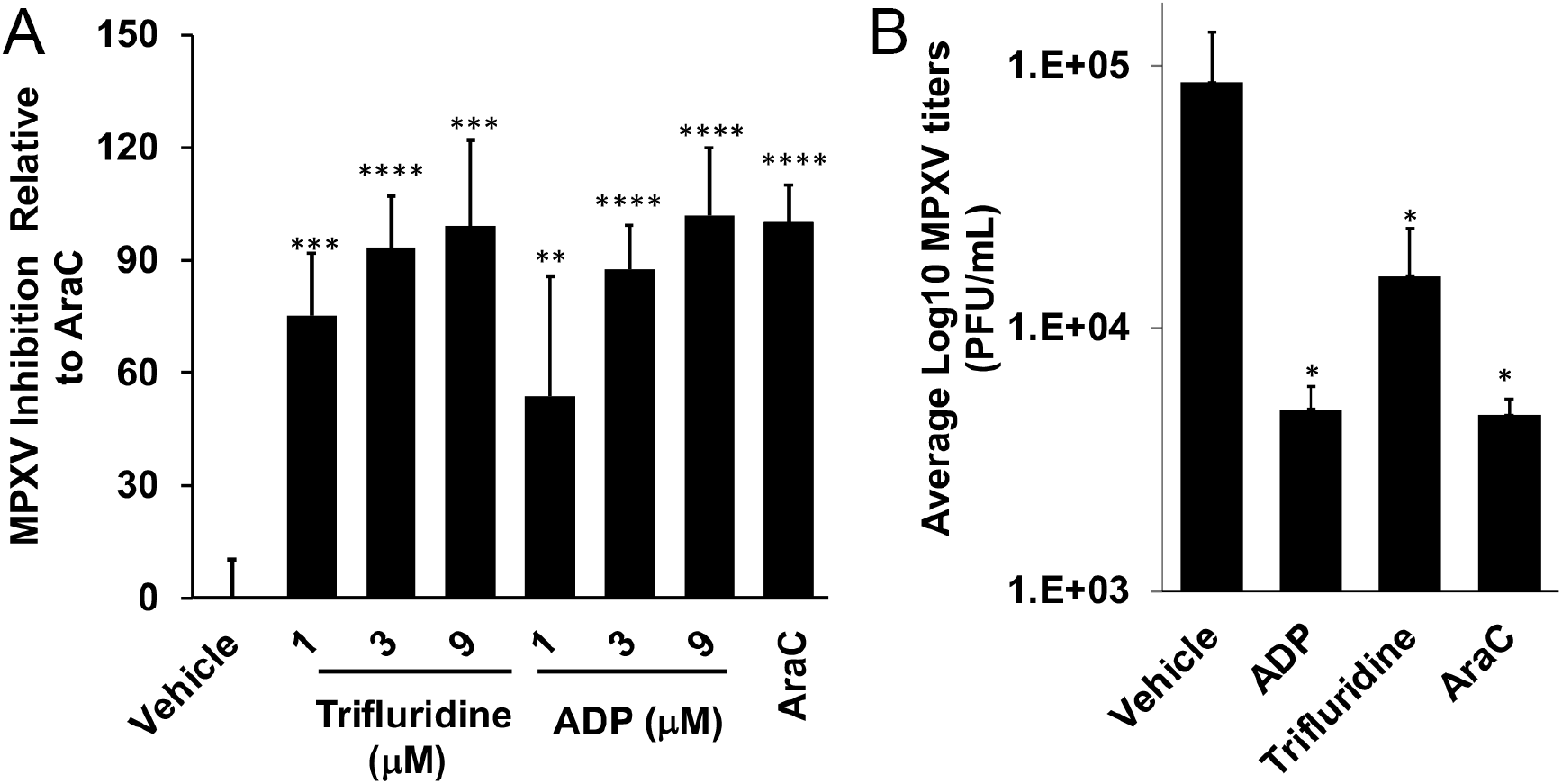
Inhibition of MPXV replication by trifluridine and Adefovir dipivoxil (ADP). **(A)** HFFs were infected with MPXV-WA-2003-Fluc (luc+MPXV) under an early/late promoter (MOI=0.01) treated with indicated compounds at indicated concentration for 24 h. Firefly luciferase activities were measured. The inhibition by AraC were normalized to 100. **(B)** MPXV-MA001 2022 isolate was added to cells at an MOI of 2 for 1 h. Virus was removed, cells were washed with PBS and trifluridine or adefovir dipivoxil was added at 10 µM. Cells were harvested 24 h post infection. AraC treatment was used as the positive control. Viral yields were titrated using a plaque assay on E6 cells. Each treatment was tested in triplicate. *p ≤ 0.05, **0.001<p ≤ 0.01, ***0.0001<p ≤ 0.001, ****0.00001<p ≤ 0.0001, ns, not significant.

## Discussion

Nucleos(t)ide analogs comprise a main class of antiviral drugs^37,38^, as exemplified by various herpes virus inhibitors^39^, a large panel of nucleos(t)ide reverse transcriptase inhibitors against HIV^40^ and / or hepatitis B virus (HBV)^41^, hepatitis C virus (HCV) inhibitors^42^, and recently, SARS-CoV-2^43-46^. Mechanistically, nucleoside analogs are intracellularly converted to the active triphosphate (TP) form, sequentially via monophosphate (MP) and diphosphate (DP), by host or virally-encoded kinases. The TPs then compete against endogenous nucleoside triphosphates (NTPs) for incorporation by the viral polymerase. Once incorporated, these analogs act as chain terminators to stall viral genome replication. In cases where the intracellular conversion into MP, typically the rate-limiting step of nucleoside drug bioactivation, is insufficient, the MP is chemically installed to bypass kinase functions, constituting a mechanistically distinct nucleotide drug family. The FDA-approved smallpox drug BCV is a prodrug of nucleotide drug cidofovir, which belongs to the acyclic nucleoside phosphonate (ANP)^47^ sub-class. Important antiviral drugs of this sub-class also include reverse transcriptase inhibitors tenofovir^48,49^ for treating HIV and HBV, and adefovir^50,51^, an HBV drug, typically administered in an ester prodrug form to overcome the low cell permeability. The two drugs characterized in this study, trifluridine and adefovir dipivoxil represent these two pharmacologically distinct classes of nucleos(t)ide drugs.

Trifluridine, or 5-trifluromethyl-2′-deoxyuridine, is a classic nucleoside drug which has long been used to treat herpes simplex virus (HSV) infection of the eyes (keratoconjunctivitis)^52^. Trifluridine was previously shown to have anti-VACV activity and used to treat VACV infection of eyes ^53,54^, although its potency was nunclear. Interestingly, trifluridine was used to treat the eye complications of mpox in the 2022 outbreak ^55^. In addition, trifluridine is also approved for treating colorectal cancer and gastric cancer. Here our data showed its low nM potency on VACV infection of HFFs and strong antiviral effect on MPXV replication. Mechanistically, trifluridine is intracellularly converted into the active TP form (**7, Fig. 6A**) by cellular/viral thymidine kinases. As a thymidine analog, TP **7** competes against the endogenous TTP for incorporation by the viral DNA polymerase. The incorporated trifluridine terminates DNA by virtue of its 5-trifluoromethyl group on the uracil base which disrupts proper base paring (**Fig. 6A**). However, trifluridine is labile toward degradation by thymidine phosphorylase, and thus requires the use of a thymidine phosphorylase inhibitor tipiracil in a combination setting for systemic cancer treatment^56^. The other drug studies herein is adefovir dipivoxil^51^, a prodrug of ANP adefovir for HBV treatment. Upon cellular uptake, adefovir dipivoxil is cleaved by a cellular carboxylesterase (CES) to release the ester promoiety and generate Adefovir (**8, Fig. 6B**). This is followed by two successive phosphorylation by AMP kinase to produce the active adefovir-DP^50^ (**9, Fig. 6B**). By competing against cellular dATP, adefovir-DP is incorporated by the viral DNA polymerase, and subsequently causes an obligate chain termination due to the lack of the 3′OH group. Against VACV and MPXV replication in HFFs, both trifluridine and adefovir dipivoxil showed potent inhibition with EC_50_s in the nM range, without discernable cytotoxicity (CC_50_ > 250 µM). These results validate both drugs as viable candidates to be further investigated as potential anti-MPXV drugs. The successful repurposing of adefovir dipivoxil will add to the already approved BCV to further enhance ANP prodrugs as an important drug class for treating poxvirus infections. In addition, nucleoside analog trifluridine as a poxvirus drug candidate will introduce a mechanistically distinct drug class and expand the options for synergistic combination therapies with ANPs or tecovirimat.

**Fig. 6.**
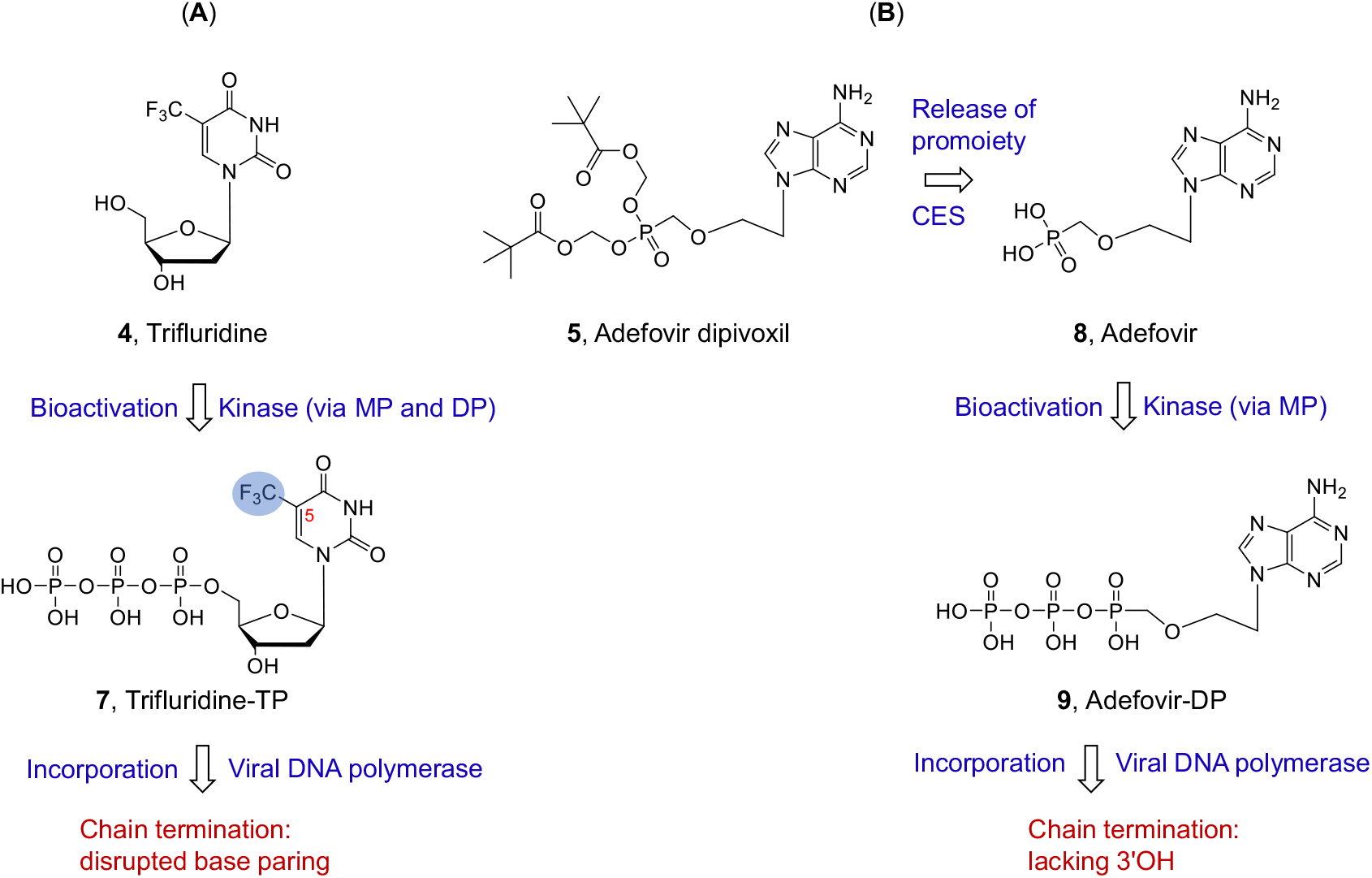
Mechanism of action of nucleoside analog trifluridine (**4**) and nucleotide analog prodrug adefovir dipivoxil (**5**). **(A)** Trifluridine is intracellularly converted into TP by thymidine kinases. Upon incorporation by viral DNA polymerase, the -CF_3_ group at the 5 position disrupts base paring and terminates viral DNA. **(B)** Ester prodrug adefovir dipivoxil is intracellularly converted into adefovir first under the action of carboxylesterase (CES). The subsequent phosphorylation by AMP kinase produces the active adefovir-DP. When incorporated, obligate termination of the viral DNA chain occurs due to the lack of the 3′OH.

Here in this study we also validated the utility of the Gluc expression VACV under a late promoter F17R (vLGluc) in measuring EC_50_ of compounds in poxvirus drug research. Because Gluc is secreted into the medium ^57^, this assay is rapid, non-disruptive, and highly simplified VACV replication reporter with exceptionally high Signal-to-Basal ratio by measuring Gluc activities in the media ^28^. This reporter VACV was suitable for high-throughput screening as shown in our previous study ^28^, which will further facilitate our future research to identify poxvirus inhibitors.

In summary, we characterized the antiviral activities of nucleoside analog trifluridine and ANP adefovir dipivoxil against VACV and mpox in primary fibroblasts. Further testing in animal models would inform their *in vivo* anti-MPXV activities. Chemical modification of the compounds may also improve their potency, pharmacokinetic (PK) and safety profiles.

## Materials and Methods

### Viruses and cells

Human Foreskin Fibroblasts (HFFs) were obtained from Dr. Nicholas Wallace at Kansas State University. HFFs and E6 cells (ATCC-CRL-1586) were cultured in Dulbecco’s minimal essential medium (DMEM; Fisher Scientific). BS-C-1 cells (ATCC-CCL26) were cultured in Eagle’s Minimum Essential Medium (EMEM). The EMEM or DMEM were supplemented with 10% fetal bovine serum (FBS: VWR), L-glutamine (2 mM, VWR), streptomycin (100 μg/mL, VWR), and penicillin (100 units/mL, VWR). Cells were cultured in an incubator with 5% CO_2_ at 37°C.

Vaccinia virus Western Reserve (WR, ATCC VR-1354) strain was propagated and purified by sucrose cushion as described previously ^58^. MPXV-WA 2003-044 ^59^ and an MPXV-MA001 2022 isolate (GenBank: ON563414.3) were utilized in this study. Recombinant VACV expressing Gaussia luciferase under VACV early, intermediate, or late promoter vEGluc, vIGluc, and vLGluc were described previously ^30^. Preparation and infection of VACV and MPXV were carried out as described previously ^60,61^.

### Titration of VACV and MPXV by plaque assay

Titration of VACV and MPXV by plaque assay were carried out as described previously ^60^. BS-C-1(for VACV) or E6 (for MPXV) cells were cultured in 6- or 12-well plates and infected with diluted virus samples and incubated in culture medium (VACV, EMEM, 2.5% FBS; MPXV, DMEM, 2% FBS) (for MPXV) and 0.5-1% methyl cellulose for 48 h (VACV) or 96 h (MPXV). Cells were stained with 0.1% crystal violet for 5 min, followed by washing with water and the number of plaques were counted.

### Chemicals

Cytarabine (AraC) was purchased from Sigma-adlrich. Brincidofovir (BCV), trifluridine and adefovir dipivoxil were purchased from TargetMol.

### Cell Viability Assay

Cell viability was measured by typan blue staining or MTT assay. For typan blue staining assay, cells were cultured in the presence of DMSO or specific compound at a desired concentration. Cells were then examined using trypan-blue exclusion as described elsewhere ^62^. MTT (3-(4,5-dimethylthiazol-2-yl)-2,5 diphenyl tetrazolium bromide) assay was performed using an MTT assay kit (Cayman Chemical). Cells in 96-well plates were treated with DMSO or desired chemical inhibitors at different concentrations and incubated for desired time. Ten µL of MTT reagent were added to each well, and cells were incubated for 3 h. A 100 µL crystal resolving solution was added to each well, and absorbance at 595 nm was measured using the citation 5 imaging reader (UV light) was measured for each well after 18 h of incubation at 37 °C.

### Determination of half maximal effective concentration (EC_50_)

HFFs were cultured in 96-well plates. The cells were infected with vLGluc at a multiplicity of infection (MOI) of 0.01 in the presence of DMSO or specific compound at a series of concentrations. Gluc activities were measured 24 hpi. The EC_50_ was calculated using the equation: log(inhibitor) vs. normalized response – variable slope in GraphPad Prism software (version 9.5.0).

### Luciferase Assay

Gaussia luciferase activities in culture medium were measured using a Pierce Gaussia luciferase flash assay kit (Thermo Scientific) using a GloMax luminometer according to manufacturer’s instructions.

Firefly luciferase activities for MPXV experiments were measured using an ENSPIRE plate reader (PerkinElmer, Waltham, MA) using the Luciferase Assay System (Promega, Madison, WI, United States) according to manufacturer’s instructions.

### Quantitative Real-Time PCR

Total DNA was extracted using E.Z.N.A. Blood DNA Kit. Relative viral DNA levels were quantified by CFX96 real-time PCR instrument (Bio-Rad, Hercules, CA) using All-in-oneTM 2× qPCR mix (GeneCopoeia) with specific VACV primers against the C11 gene. 18S rRNA gene primers were used as the internal reference.

### Statistical Analysis

All data were represented as the means of at least three independent experiments. Student’s T-test was used to access for significant difference between two means with P ≤ 0.05.

## Acknowledgment

We thank Nicholas Wallace (Kansas State University) for providing HFFs. We thank Bernard Moss for providing VACV.

Z.Y. was supported in part by grants from the National Institutes of Health (R01AI128406).

P.S.S is supported in part by grants from the Biomedical Advanced Research

The findings and conclusions in this report are those of the authors and do not necessarily represent the official position of the Centers for Disease Control and Prevention.

